# Circadian regulation of microRNA-target chimeras in *Drosophila*

**DOI:** 10.1101/622183

**Authors:** Xiju Xia, Xiaonan Fu, Binbin Wu, Jinsong Zhu, Zhangwu Zhao

## Abstract

MicroRNA is critical coordinator to circadian regulation by silencing gene expression. Although many circadian related miRNAs and some of its target are known, the global functional miRNA-mRNA interaction networks remain poorly understand which is hindered by imperfect base-pairing between miRNA and target mRNA. In this study, we used CLEAR (Covalent Ligation of Endogenous Argonaute-bound RNAs) -CLIP (Cross-Linking and Immuno-Precipitation) to explore the regulatory functions of miRNAs in the circadian system by comparing the miRNA-mRNA interactions between the *Drosophila* wild-type strain *w^1118^* and the *Clk* mutant *Clk^jrk^*. We unambiguously identified thousands of miRNA-mRNA interactions from CLEAR-CLIP data set at unprecedented depth *in vivo* for the first time. Among them, about 300 miRNA-mRNA interactions were involved in the regulation of circadian, in which miRNAs targeting core clock genes *pdp1*, *tim* and *vri* presented distinct changes in response to *Clk^jrk^*. Particularly, the *mir-375-timeless* interaction from CLER-CLIP shows important effects on circadian, this functional event occurred in the l-LNv neurons. Overexpression of *mir-375* in *tim* neurons caused decreases in TIM content resulting in arrhythmicity of daily locomotion and changes of sleep. This present work provides a global view of miRNA targeting in the circadian rhythm.

## Introduction

In animals, the intrinsic circadian clock regulates daily rhythms in physiology and behavior, which are entrained by environmental stimuli such as light and temperature^1, 2^. This robust timing system maintains rhythmic oscillation even under constant darkness conditions in the fruit fly *Drosophila melanogaster*.

Sleep is one of the established circadian behavior, which is highly associated with health status^3^. Sleep in *Drosophila* has been well characterized by certain parameters such as total sleep, sleep bout duration and sleep bout number^4–6^. Throughout the daily sleep-wake cycle, the flies exhibit two peaks of activity - one before lights on and one before lights off. Genetic dissection in *Drosophila* indicates that sleep is regulated by the circadian clock. The mutations of the core clock genes result in abnormal sleep^7, 8^, while the neuropeptide pigment dispersing factor (PDF)-expressing peptidergic clock neurons regulate arousal as well as sleep stability^9^.

The *Drosophila* rhythmic behavior is maintained by a central clock network in the brain, which is consisted of ~150 circadian neurons^10^. These clock neurons include ventral lateral neurons (LNvs) (I-LNvs, s-LNvs, and 5^th^ s-LNvs), dorsal lateral neurons (LNds), lateral posterior neurons (LPN) and dorsal neurons (DN1, DN2, and DN3), classified based on anatomical locations and varied expression of core clock genes^10–13^. Among them, the PDF-positive neurons (I-LNvs and s-LNvs) are essential for normal circadian activity and daily sleep. Their ablation results in arrhythmic flies^4^.

The circadian clocks are modulated by a complex transcriptional and translational feedback loop (TTFL) in both mammals and flies^5–7^. In *D. melanogaster*, two interlocked clock feedback loops are known to regulate the circadian rhythm. They consist of *period* (*per*), *timeless* (*tim*), *clock* (*clk*), *cycle* (*cyc*), *vrille* (*vri*) and *par domain protein 1e* (*PDP1e*). The transcription factors CLK and CYC form a heterodimer that binds to the E-box of *per* and *tim* promoters, thereby promoting their transcription^8, 9^. The cycling expression of *tim* mRNA with a peak at Zeitgeber time (ZT) 14 and a trough at ZT 2 is vital for circadian output^10, 11^. Another feedback loop consists of the *clk, cyc, vri* and *PDP1e*^8, 9, 12–14^. However, the mechanism of how these feedback loops sustain ~24 h rhythm is still unclear, since multi-layered regulation involving post-transcriptional and post-translational regulation are imposed to produce the daily rhythm^15–17^.

The highly conserved and widespread microRNAs (miRNAs) play a critical role in post-transcriptional gene regulation^18–22^. They are loaded into Argonaute (AGO) proteins to mediate mRNA cleavage or translational inhibition via base pairing with target mRNAs^20, 23–25^. Disruption of the miRNA biogenesis pathway considerably weakens the rhythm in locomotor activity^26^. Although the rhythmically expressed miRNAs have been identified^27–29^, only a limited set of miRNA-target pairs have been verified to affect circadian rhythms^26, 30–33^. Decoding of the miRNA-mRNA interaction network is difficult because of incomplete complementary pairing between miRNAs and targets as well as context-dependent dynamics of the miRNA-target interactions. As a result, some miRNAs such as *miR-124* and *miR-959-964* have been shown to be involved in circadian rhythms but without known targets^29, 34, 35^. Thus, a global identification of circadian-related miRNA-target interactions will greatly improve our understanding of the role of miRNAs in circadian regulation.

The effort to accurately predict the miRNA-targets is compromised by the complex and dynamic cellular interaction networks^36, 37^. Immunoprecipitation of the miRNA effector protein Argonaute followed by microarray^26^ or RNA-seq analysis^38^ narrows down the search scope of potential targets but still relies on bioinformatic analysis to predict miRNA-mRNA pairing. The recently improved experimental method of the CLEAR (Covalent Ligation of Endogenous Argonaute-bound RNAs)–CLIP, which isolates miRNA-mRNA chimeras from endogenous AGO-miRNA-mRNA complexes, permits unambiguous identification of miRNA’s targets, which provide a snapshot of true, physiological miRNA-mRNA interactions *in vivo*^24, 39–41^.

To systematically investigate the role of miRNAs in circadian rhythms, we used the CLEAR-CLIP assay in *Drosophila*. Tens of thousands of miRNA-mRNA interactions were detected highlighting the well-established miRNAs targeting features. Our data indicate that miRNAs extensively fine-tune the expression of circadian-relevant genes. Importantly, we show here that the regulation of *tim* by *miR-375* plays an important role in locomotor activity and sleep. This present work provides a global view of the miRNA-mRNA interactions in circadian rhythms and a solid foundation for future functional studies.

## Results

### Identification of miRNA-mRNA interactions by CLEAR-CLIP

The brain clock neurons are the regulatory center of circadian rhythm^42^, in which the transcription factor *Clk* is essentially required for rhythmic behavior. Ablation of *Clk* expression in the mutant *Clk^jrk^* makes the oscillation of miRNAs disappear in post-transcriptional regulation^29^. To uncover the physiological miRNA-mRNA interactions contributing to the circadian rhythm, we applied a modified Ago1 CLEAR-CLIP to separately analyze the head and the whole body in *Clk^jrk^* and *w^1118^* flies (Figure S1). The purified Ago1-miRNA-mRNA complexes could be either self-ligated as chimeric reads or remain unchanged mapping to a single locus (CLIP peaks) (Fig. 1A). In total, we obtained 213,404,300 reads from 8 libraries (Table S1A), of which 88.32% could be mapped to mRNAs, and 1.92% from miRNA-target chimeras (Fig. 1B). The recovery of previously experimentally supported miRNA-mRNA interactions in *Drosophila* showed a reliable dataset (Table S1B), exemplified by the verified regulation of hid by bantam^43^ (Fig. 1A).

**Fig. 1.**
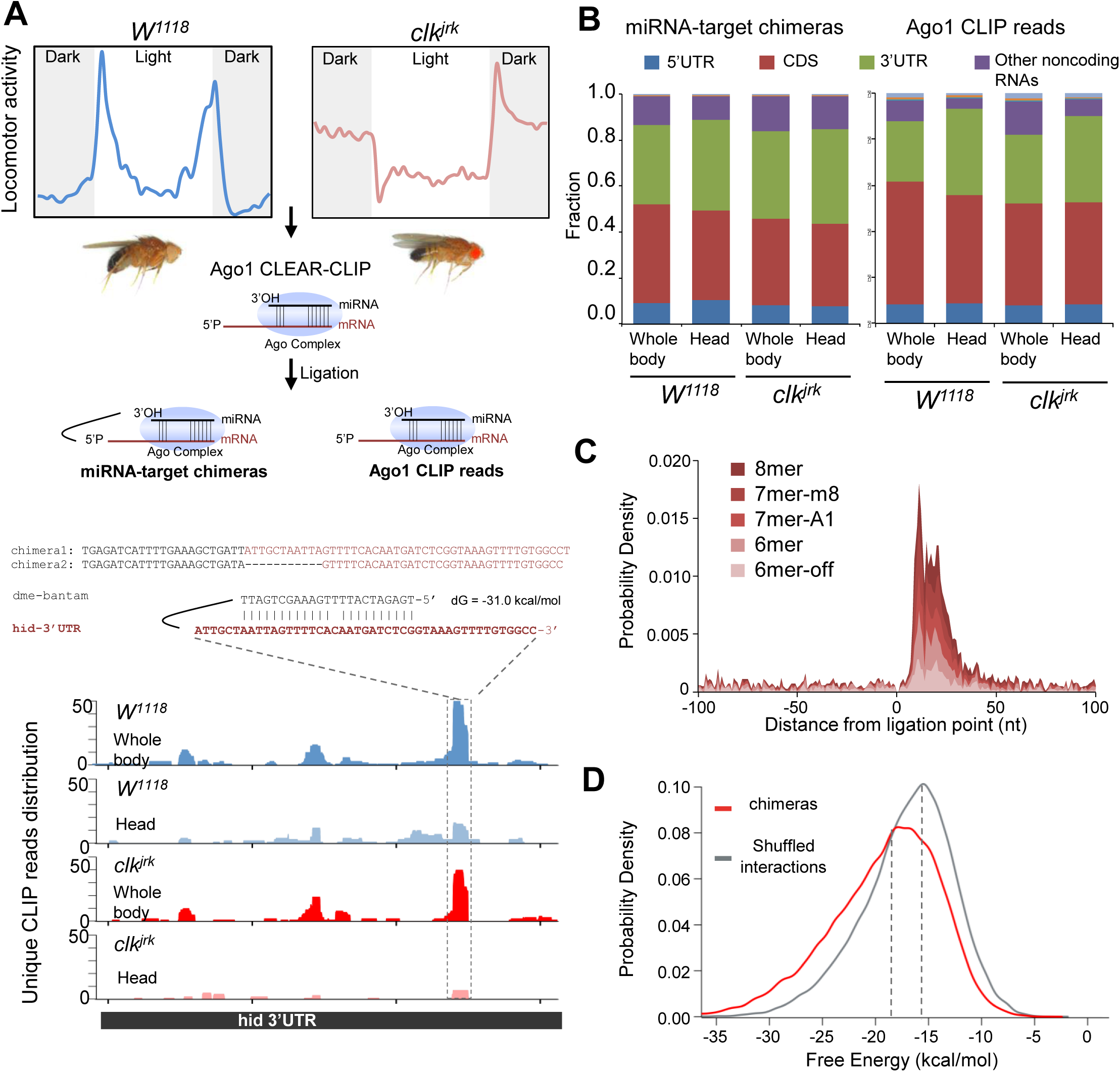
Ago1 CLEAR-CLIP decodes miRNA-mRNA interactions in Drosophila. (A) Schematics of CLEAR-CLIP to detect miRNA-mRNA interactions in Drosophila. Wild-type *W^1118^* and Clk mutant *clk^jrk^* fiies with arrhythmic phenotype were collected. The UV-crosslinked sample was homogenized in lysis buffer. Ago1-RNA complex was immunoprecipitated and incubated with T4 RNA ligase to favor the direct ligation of miRNA with its target mRNA fragments. The recovered RNAs were subjected to library construction and sequencing. With the bioinformatics analysis, the chimeric reads could be identified along with the Ago1 binding reads, exemplified by the known bantam targeting hid. (B) The fraction of different RNA classes annotated to Drosophila for miRNA-target chimeras and Ago CLIP reads. (C) Density plot of canonical miRNA seed matches in target regions relative to ligation site for all the chimeras. (D) The median predicted binding energy between miRNA and matching target mRNA found in chimeras was stronger by over 2.8 kcal mol−1 than in randomly matched pairs.

In this study, we discovered 15,825 miRNA-target interactions in total involving 239 mature miRNAs and 6,540 mRNA transcripts in *Drosophila* (Table S2A, B). These miRNA-mRNA reads were further classified to miR-first and miR-last chimeras according to the location of miRNA (Table S1C). The miR-first chimeras (92.55%) were predominant compared with the miR-last chimeras (7.45%), and they showed a higher correlation coefficient between miRNA frequency and miRNA abundance (r≥0.79), while the correlation coefficient was low (r =~0.45) for miR-last chimeras (Fig S2A, B). Thus, we focused exclusively on the miR-first chimeras in subsequent data analysis.

The target sequences from the miR-first chimeras were enriched with canonical seed which matched within 50 nt of the ligation sites (Fig. 1C). In addition, the mean of predicted free energy between miRNAs and the matched target mRNAs found in chimeras was lower by 2.8 kcal mol^−1^ than that in randomly matched pairs (Fig. 1D, p<0.001). The strong binding energies of chimeric reads suggest that stable chimera stems come from genuine mRNA-miRNA pairing in miRISC rather than from proximity-induced ligation of non-specific RNAs in the solution. These data indicated that the CLEAR-CLIP chimeras reveal a reliable and high-resolution miRNA-target interaction map.

### The miRNA-target chimeras preserve functional interaction

To further evaluate the physiological functions of miRNA-target interactions, we analyzed the binding motif enrichment of *let-7-5p*^44^, *miR-34-5p*^45^, *miR-305-5p^46^* and *miR-276a*^47^, which have been investigated in circadian regulation. The enriched 7-mer motifs in the targets of these miRNA were reverse-complementary to the miRNA seed regions, indicating the crucial role of the seed sequences in miRNA targeting (see Fig. 2A and Fig. S2C). To further examine the evolutionary conservation of chimera-defined target sites, the PhyloP conservation scores of 3’UTR targeting sites were retrieved from multiple alignments of 27 insect genomes. We found that the conservation score of chimera-defined targets displayed a marked increase at 6-nt interacted stem relative to flanking regions (Fig. 2B), indicating the close physiological relevance.

**Fig. 2.**
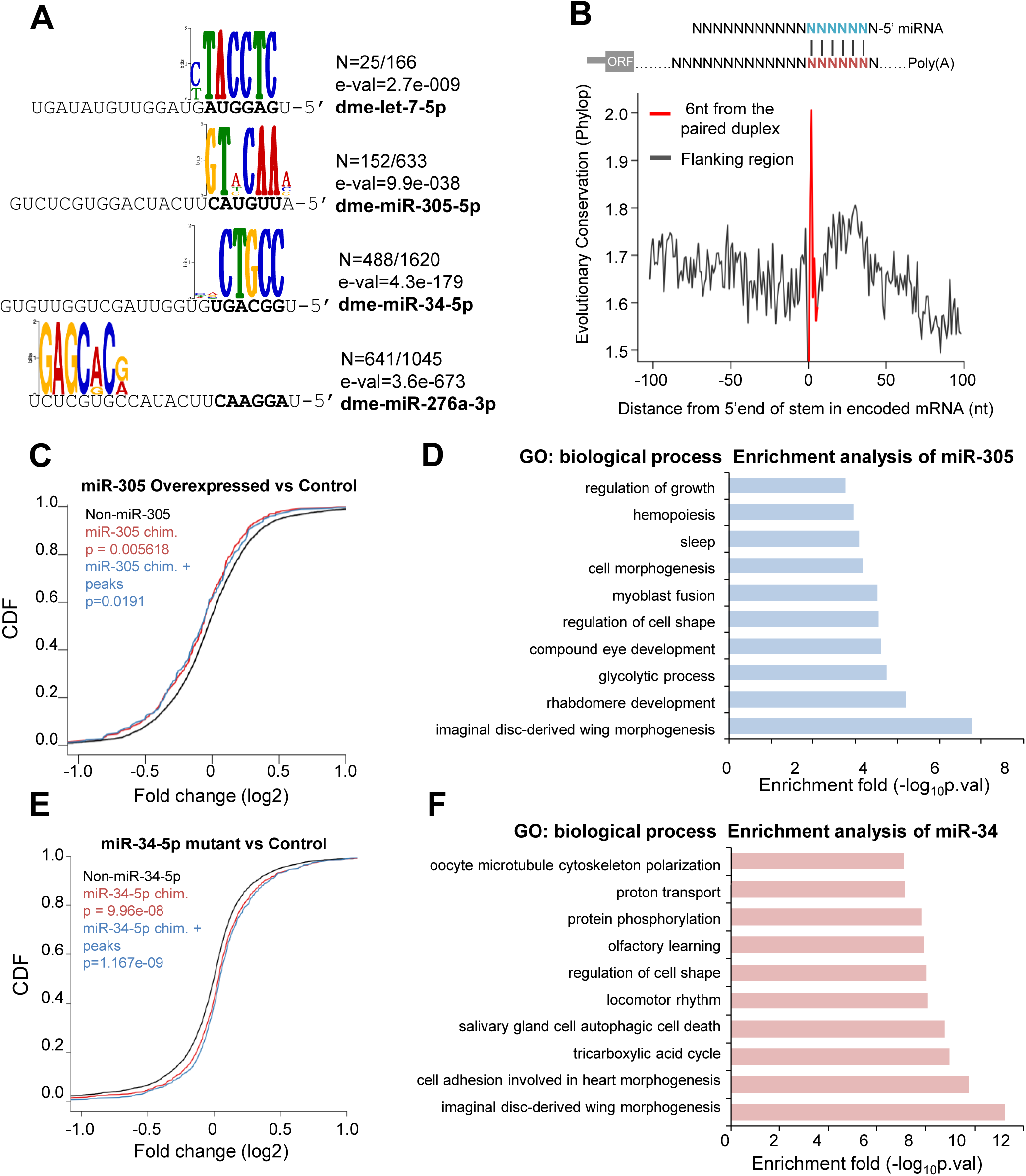
miRNA-target chimeras preserve physiological relevant interactions. (A) Enriched miRNA binding motifs by chimeras-defined target sites. n, number of motifs found/total number of targets analyzed. E-val, e-value of the motif returned by MEME. Most motifs are complementary to the miRNA seed (bold). (B) Average conservation score of perfect seed matches from miRNA:target interactions located in 3’UTR defined by chimeras. The Phylop conservation score for 27 insect species was downloaded from UCSC. (C) Cumulative distribution function (CDF) plot shows log_2_-transformed expression differences for mRNAs identified in miR-305 chimeras comparing the other mRNAs which is not defined as miR-305 targets. The analysis was performed using the transcriptome measurement in miR-305 overexpressed fly versus the wild type control. *P*-values was calculated from Kolmogorov–Smirnov testing. (E) Cumulative distribution function (CDF) plot shows the change in transcript level after knocking out miR-34-5p. The similar analysis as (A) presents the mRNA expression distribution detected in miR-34-5p chimeras, miR-34-5p chimeras supported by Ago1 peaks and the other mRNAs. (D,F) GO enrichment analysis of chimeras-defined miRNA (E) targets. The top ten biological process terms were plotted.

In addition to assessing the functional outcome of the chimera-identified miRNA-target by CLEAR-CLIP, we also analyzed how the altered RNA levels of *miR-305* and *miR-34* affected the expression of their target mRNAs based on previously published data related to rhythmicity^45, 46^. In the CLEAR-CLIP dataset, the *miR-305* bound to 598 target sites within 555 mRNA transcripts. Overexpression of *miR-305* in *Drosophila* considerably lowers the abundance of mRNAs bearing chimera-defined *miR-305* target sites, compared with the mRNAs that had no *miR-305* target site (p=0.005618). And the subset of these mRNAs supported by both chimeras and Ago1 CLIP peaks was also significantly repressed (p=0.0191) (Fig.2C). Functional enrichment analysis of *miR-305* targets also demonstrated their important roles in sleep (Fig. 2D), which has been observed in the *miR-305* mutant strain^46^. Furthermore, the mRNA levels of the *miR-34* targets were significantly elevated in the *miR-34* null mutant (*miR-34-5p*), compared with *w^1118^* (p<0.001) (Fig. 2E). In this study, GO enrichment analysis of the chimera-defined *miR-34* targets linked *miR-34* to aging and brain disease (Fig.2F), consistent with the previous report that *miR-34* is associated with aging and neurodegeneration^45^. These results verified the physiological significance of the miRNA-mRNA interactions identified by CLEAR-CLIP.

### *Clk*-oriented miRNA-mRNA interactions are critical for circadian rhythm

To explore the whole miRNA-mRNA interactions network in the circadian system, we visualized 9325 unique miRNA-target chimeras supported by corresponding Ago1 CLIP peaks with SOM clustering (Fig. S3A). The miRNA-mRNA interactions were significantly changed in *Clk^jrk^* (Fig.3A), implying widespread circadian impact by CLK. Furthermore, by quantitatively identifying the affected miRNA-mRNA interactome (fold change>2), we found that 53.0% of the circadian relevant regulations were altered, respectively (Fig. 3A). For example, the interaction between *miR-276* and *tim* only appeared in the wildtype strain *w^1118^*. But the *miR-34*- *pdp1* pairing only happened within the mutant strain *Clk^jrk^.* These phenomena implicate that the miRNAs involved in circadian regulation are dynamically adapting to the systemic changes.

To understand the functional impact of changed miRNA-mRNA interactions, we performed a functional enrichment analysis separately using the significantly up/down-regulated genes. Results showed that many circadian-related functional groups such as sleep and visual perception were top-ranked, especially enriched in the fly-head groups, while chemical synaptic transmission is notably affected after *Clk* mutation (Fig. S3C).

Moreover, by grouping the genes involved in circadian phenotype, we found that 299 miRNA-mRNA interactions involving 9 core circadian genes were plotted as a miRNA-regulated network (Fig. S3B), in which a majority of interactions appeared to target the three *Clk*-downstream genes *pdp1*, *vri* and *tim* (Fig. 3B). To compare the miRNA-involved regulations in these three genes, we retrieved the interaction pattern under different conditions. Results displayed distinct regulatory features in response to *Clk* disruption (Fig. 3C). The miRNAs targeting *pdp1* acted in a *Clk*-independent manner. Conversely, the interactions between *tim* and miRNAs occurred in *w^1118^* but were not detectable in *Clk^jrk^*, with exceptions of *miR-193-3p* and *miR-2c-3p*. Binding of miRNAs to the *vri* transcript exhibited a mixed pattern of both *Clk*-dependence and *Clk*-independence. The *miR-276a/b-3p*, *miR-14-3p*, and *miR-210-3p* were joint regulators of these three genes (Fig. 3D). Alternatively, *miR-375-3p* and *miR-305-5p* only targeted *tim* in wild type flies (Fig. S4B), in which *miR-305-5p* had been reported to target *tim* in several circadian screens^48–50^. These findings suggest that gene regulation by miRNAs has a profound impact on circadian system stability.

### Oscillations of *Mir-375* expression in pacemaker neurons is affected in *Clk^jrk^*

Besides the reads mapped to mRNA, the CLEAR-CLIP dataset also included lots of unligated miRNAs, which has been suggested as a reliable indicator of functional miRNA abundance^51^. To understand the Ago1-bound miRNA changes, we summarized the reads mapped to mature miRNAs. Results showed that only a few miRNAs (*miR-375-3p*, *miR-193-3p*, *miR-133-3p*) related to core circadian genes showed significant changes between *w^1118^* and *Clk^jrk^*. Among them, *miR-375-3p* displayed the largest change, with a 3.28-fold increase detected in the fly head (Fig. 4A). Thus, we focused on the *miR-375-3p* for further study.

**Fig. 3.**
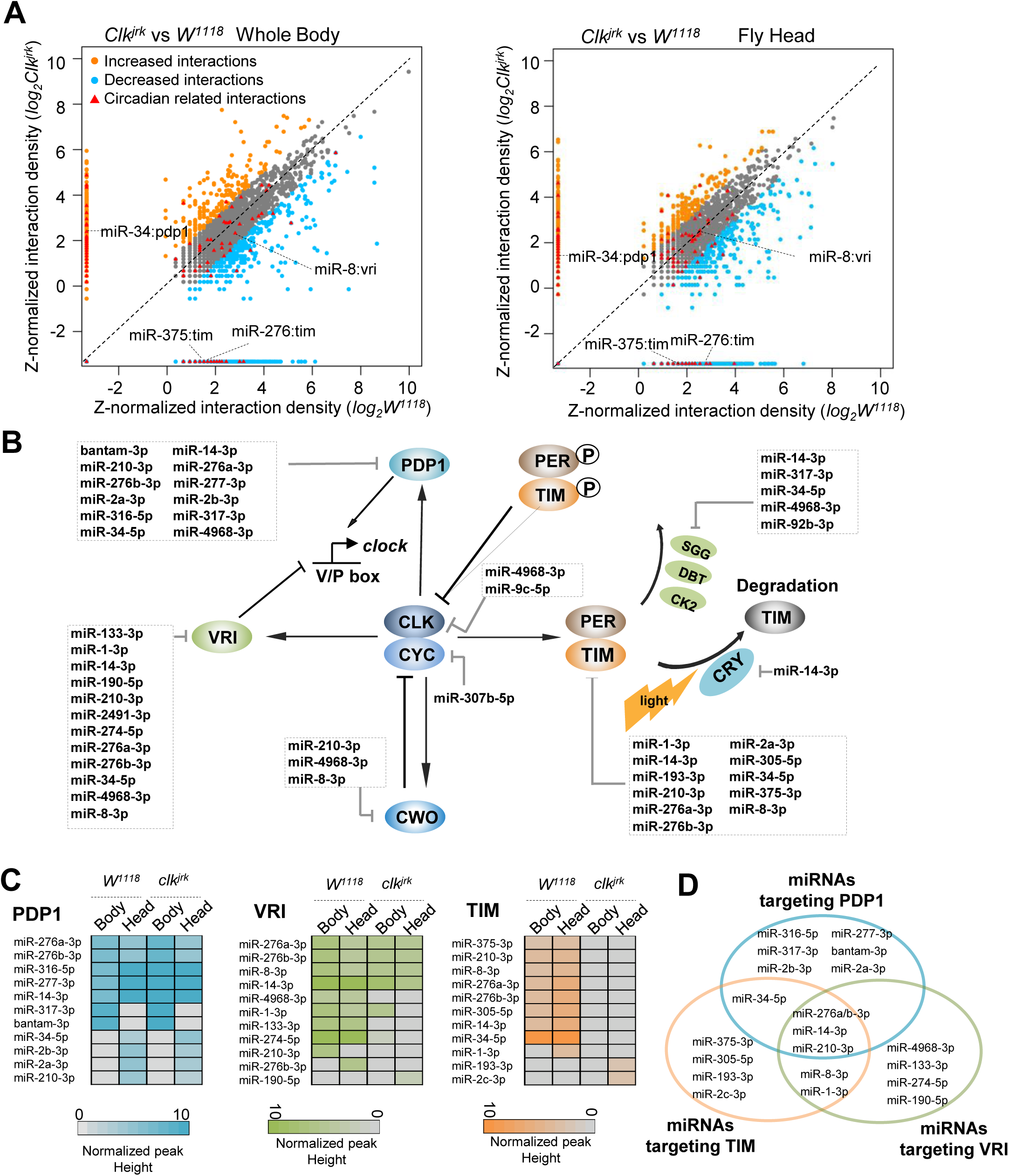
miRNA-mRNA interactions regulates circadian rhythm in Drosophila. (A) Scatter plot shows the change of miRNA-mRNA interaction comparing *Clk^jrk^* with *W^1118^*. The interactions with 2 fold changes are considered as significance. (B) The miRNAs participated regulation in the core network of circadian system. All the miRNA-mRNA interactions were supported by Ago1 peaks. (C) The heatmap shows the change of miRNA-mRNA interactions for PDP1, TIM and VRI at each condition. (D) Venn diagram shows the subsets of miRNAs contribute to the regulation of three core circadian genes.

**Fig. 4.**
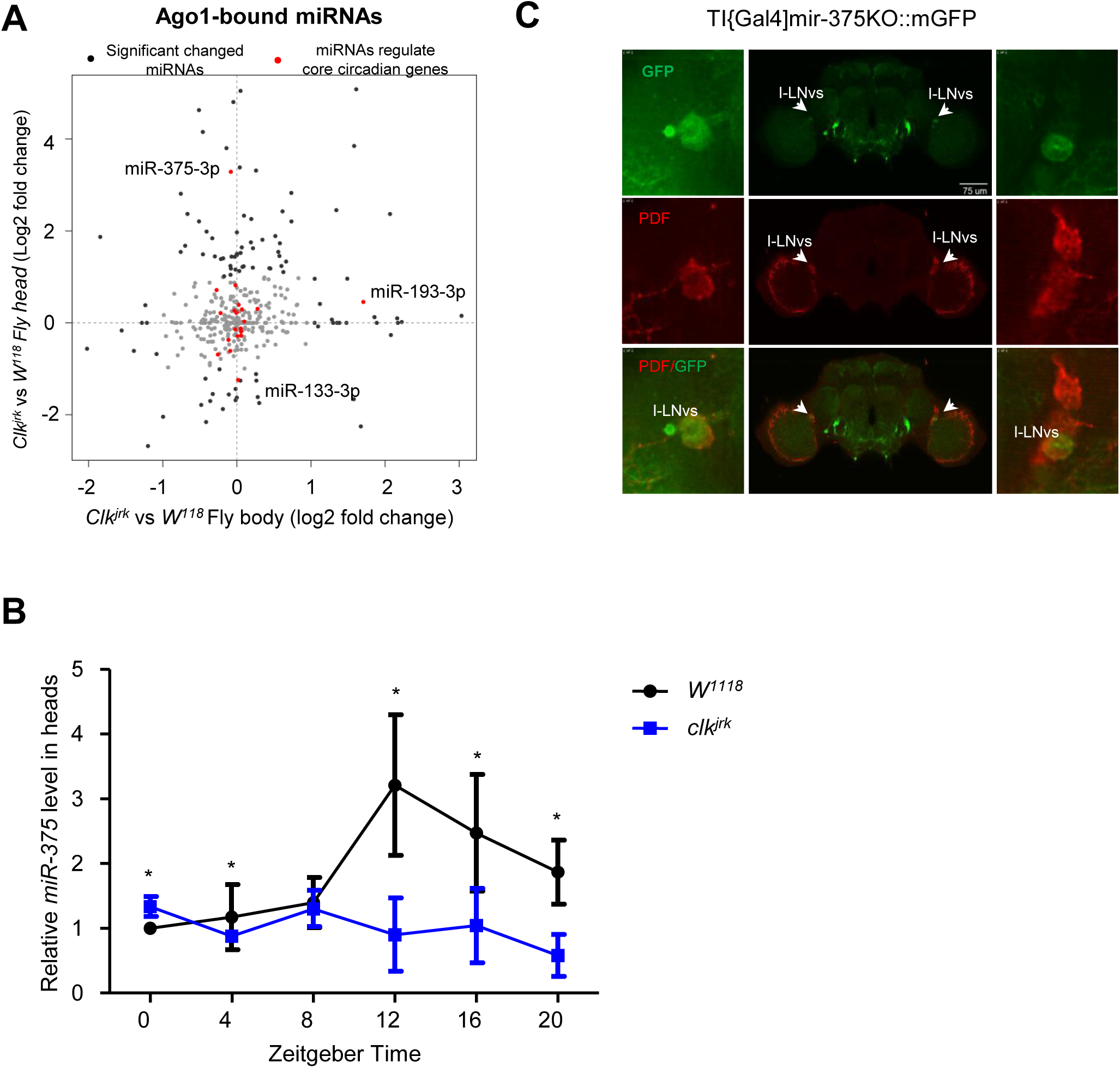
mir-375 oscillations in pacemaker neurons. (A) The changes of Ago1-bound miRNAs abundance between *Clk^jrk^* and *W^1118^* comparing whole fly and head. (B) The levels of miR-375 are altered in *Clk^jrk^* mutant flies’ heads. (C) w*; TI{GAL4}mir-375KO expression in Drosophila brain. w*; TI{GAL4}mir-375KO flies were crossed with UAS-mGFP files.

First, the daily expression of *miR-375-3p* in the fly head was analyzed by qRT-PCR at six time points (ZT0, 4, 8, 12, 16, 20) under LD condition. Results showed that it was rhythmically expressed with the peak at ZT12 in *w^1118^*, but this pattern disappeared in *Clk^jrk^* (Fig. 4B), implying its dependence on *Clk*. To investigate the spatial expression of *miR-375-3p*, we utilized a reporter strain (*TI{GAL4}mir-375[KO]* ×*UAS-Mcd8::GFP*) to monitor the expression of *mir-375*. Results showed that it was expressed in the PDF-expressing I-LNv neurons and co-localized with TIM (Fig.4C).

### *Mir-375* impacts normal circadian rhythm and sleep

To further evaluate the role of *mir-375* in circadian behavior, we overexpressed *mir-375* in *tim* neurons by using *iso:tim-gal4* driver. Results showed that the flies with enhanced expression of *mir-375* in *tim* neurons lost daily locomotor rhythm under both LD and DD conditions. Both morning and evening anticipations disappeared after the overexpression of *mir-375* (Table 1, Fig.5A-C), while control flies exhibited normal behavioral rhythms (*UAS-LUC-mir-375/+*: 95.74% rhythmic, tau=24.5 h, n=47; *iso:tim-gal4/+*: 100% rhythmic, tau=24.6 h, n=75). In addition, flies with overexpressed *mir-375* showed a decrease in sleep bout duration and an increase in sleep bout number at night compared with the control (Fig.5D, E). These results demonstrate that *mir-375* is very important in the regulation of the *Drosophila* circadian system.

**Fig. 5.**
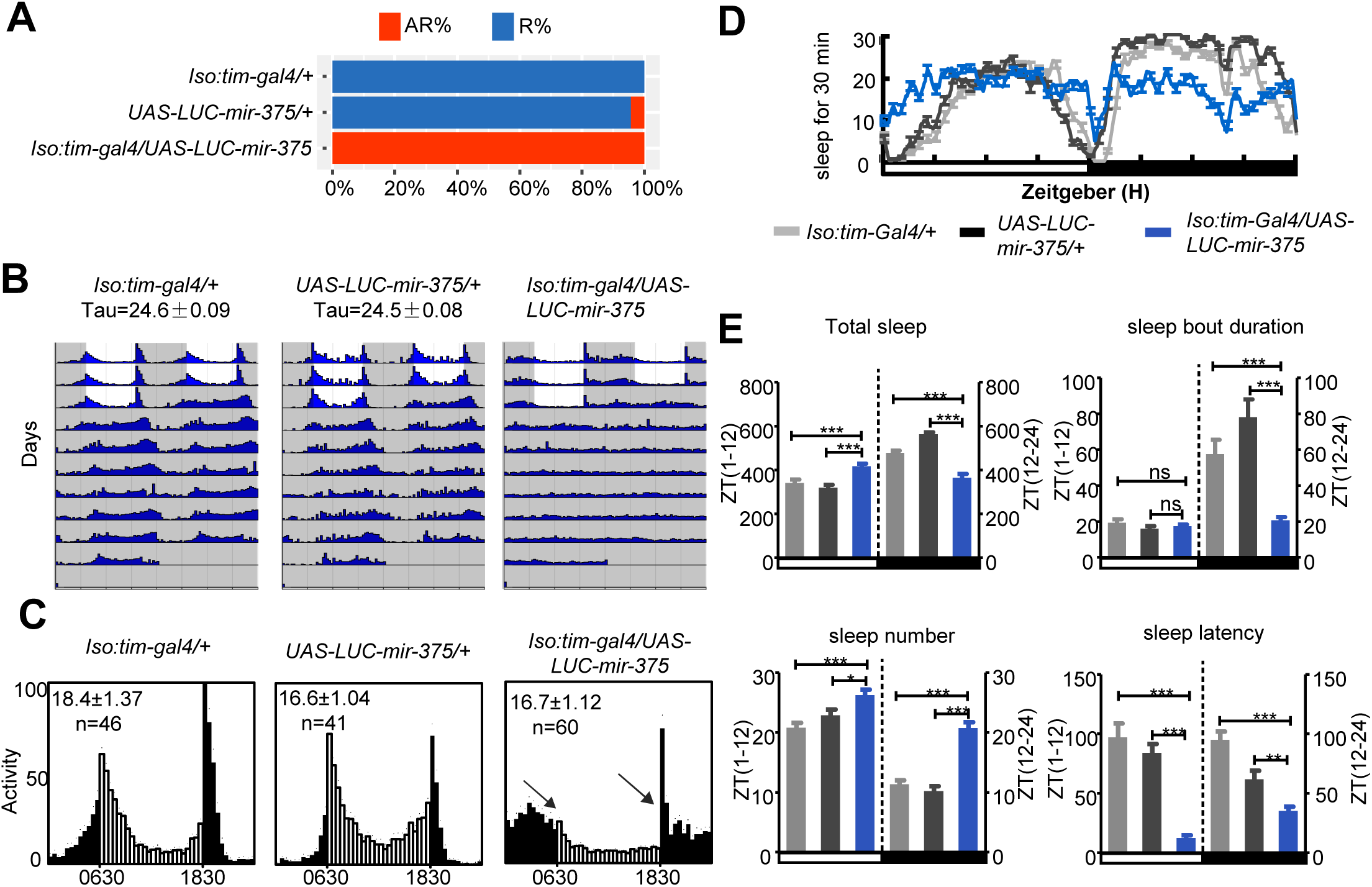
Overexpression of *mir-375* regulate circadian rhythm in *Drosophila*. (A) Overexpression of mir-375 levels causes increased arrhythmicity in flies. AR% represents the percentage of arrhythmic flies, R% represents the percentage of rhythmic flies. (B) Comparison of the circadian locomotor activity in light/dark cycles of controls and mir-375 OE flies. Histograms represent the distribution of activity through 24 h, averaged for flies over three LD days, morning and evening anticipation are lost in mir-375 OE flies. (C) Locomotor behavior under LD cycle and constant darkness. Representative double plotted actograms of iso:tim-gal4/+, UAS-LUC-mir-375/+, iso:tim-gal4/UAS-LUC-mir-375 flies. White indicates the light phase, gray indicates the dark phase. (D) Sleep analysis of male flies with overexpression of *mir-375* driven by iso-tim-gal4. The colors of gray dark, light dark and blue represent *iso:tim-gal4/+, UAS-LUC-mir-375 and iso:tim-gal4/UAS-LUC-mir-375* sleep traces. (E) Sleep profiles: total sleep, sleep bout duration, sleep number and sleep latency of male flies entrained under three LD cycles are shown, *mir-375* levels are upregulated in tim-staining neurons. Data represent mean±SEM (n = 16–32). **P<0.01, ***P< 0.001 determined by Student’s t test.

**Table 1.** Locomotor activities of flies.

| Genotype | Total flies | Rhythmic flies (%) | Period (h) | Power |
| --- | --- | --- | --- | --- |
| Iso:tim-gal4/+ | 75 | 100 | 24.6±0.09 | 56.2±6.2 |
| UAS-LUC-mir-375/+ | 47 | 95.74 | 24.5±0.08 | 65.2±7.22 |
| Iso:tim-gal4/UAS-LUC-mir-375 | 62 | 0 | * | * |
| UAS-mir-305/+ | 45 | 93.3 | 24.3±0.09 | 97.5±9.62 |
| Iso:tim-gal4/uas-luc-mir-305 | 47 | 91.5 | 23.7±0.13 | 82.8±9.33 |
| uas-luc-mir-9c/+ | 47 | 100 | 23.4±0.07 | 97.9±6.76 |
| Iso:tim-gal4/uas-luc-mir-9c | 29 | 0 | * | * |

### *mir-375* influences circadian rhythm and sleep *via* targeting *tim*

The flies with up-regulation of *mir-375* had the same phenotype as the *tim* null mutants on the regulation of the circadian rhythm (Fig.5C and Fig.S4A), and they both expressed *mir-375* in l-LNv neurons. In this study, 11 miRNA-*tim* chimera-defined target sites were founded across the *tim* transcript (Fig. S4B). Among them, the *miR-375* targeting site was located at its CDS region and the chimeras displayed a stable hybridization stem with lower free energy (Fig. 6A).

**Fig. 6.**
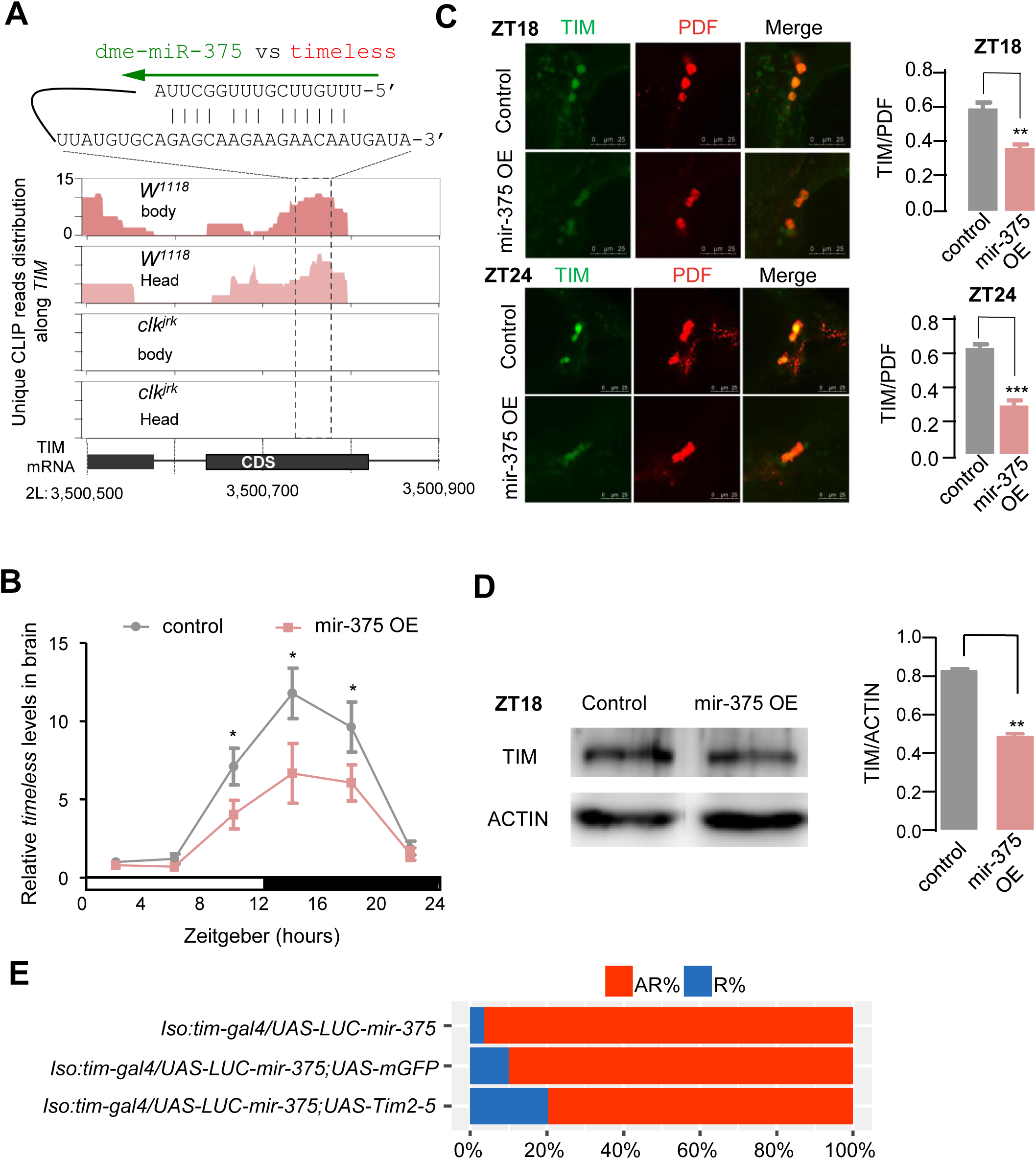
Tim is a target of *mir-375.* (A) The *Clk*-dependent interaction of miR-375 targeting timeless in its coding region. The duplex defined in chimeras was predicted by RNAhybrid. (B) Levels of *timeless* mRNA are altered in miR-375 OE flies. The relative expression levels were normalized to actin levels and were further normalized to control at ZT2. (error bar indicate SEM, \**P* < 0.05; two-tailed t test.) (C) Immunostaining of control (iso-tim-gal4/+; UAS-LUC-mir-375/+) and mir-375 OE (iso-tim-gal4/UAS-LUC-mir-375) flies against TIM (green) and PDF (red) at ZT18 (upper) and ZT24 (lower). Tim signal is normalized to the PDF signal. (n=10-17 hemisphere; error bar indicate SEM, \*\**P* < 0.01; \*\*\*\**P* < 0.0001; two-tailed t test.) (D) Western blot against TIM in control and mir-375 OE flies ZT 17, ACTIN is used as loading control, TIM levels are normalize to ACTIN levels. Protein signals are quantitated by using image J. (n=4, error bar indicate SEM, \*\**P* < 0.01; two-tailed t test.) (E) Overexpression of timeless levels recues arrhythmic behavior in mir-375OE flies. AR% represents the percentage of arrhythmic flies, R% represents the percentage of rhythmic flies.

Furthermore, to address whether *miR-375* had an effect on the expression of the core clock gene *tim*, the mRNA levels of *tim* were detected daily at six time points (ZT2, 6, 10, 14, 18, 22) in both flies with *mir-375* overexpression (OE) in *tim* neurons and control flies. Results showed that the amounts of *tim* mRNA were dramatically decreased in *mir-375* OE flies at ZT 10, 14 and 18 (Fig. 6B). Similarly, TIM protein levels in I-LNvs neurons, detected at ZT18 and ZT24 by confocal microscopy, also showed significant decreases (31.1% at ZT 18 and 54.37% at ZT24) in *mir-375* OE flies compared with control flies (Fig. 6C). This observation was also verified by western blot analysis at ZT18 (Fig. 6D). These results indicate that the TIM in the brain was significantly lower in the *mir-375* OE flies than in the control flies.

To connect arrhythmic phenotypes caused by overexpressing *miR-375* in tim-staining neurons, we simultaneously used the *UAS-tim2-5* to rescue this phenotype of overexpressing *miR-375*, in which the *UAS-*mCD8-GFP served as a control. The results showed that the *UAS-tim2-5* could reduce arrhythmicity to 80% compared with the *UAS-*mCD8-GFP control, confirming that *tim* is a real target of *miR-375* (Fig. 6E). Taken together, all these data demonstrate that the regulation of *tim* by *mir-375* plays an important role in mediating the daily circadian rhythm.

## Discussion

The significance of post-transcriptional regulation in circadian rhythms has been demonstrated by monitoring the core clock gene expression^15, 52^ and screening miRNAs involved in rhythmic behavior^44, 47–49, 53–58^. However, elucidation of the underlying molecular mechanism is impeded because only a few of the predicted miRNA-mRNA interactions have been verified. In this study, we captured miRNA-mRNA interaction pairs directly from fly tissue using the recently developed Ago1 CLEAR-CLIP^59, 60^, which enabled us to detect tens of thousands of miRNA-target interactions. Of these interactions, 75 circadian genes are regulated by 61 miRNAs in the network of 299 interactions (see Fig. 1 and Fig. S3B).

With the first global miRNA-mRNA interaction profiles in *Drosophila*, we performed multiple lines of evaluation to assess the quality of the whole dataset. First, CLEAR-CLIP recovers many known miRNA-target interactions such as *bantam:hid*^43^ and *miR-276a:tim*^47^ (Fig. 1). Second, analysis of interactions recovered from chimeras reveals general bioinformatics features for miRNA targeting, including seed enrichment, stable binding energy and evolutionary conservation (Fig.1, Fig.S3). Third, the chimera-defined target mRNAs exhibit significant changes after manipulating the corresponding miRNA (Fig. 2). Fourth, functional enrichment analysis of the targets of *miR-305* and *miR-34* are consistent with the previous phenotypic study of mutant flies^45, 46^. These together substantiate the reliability of the datasets for miRNA-mRNA interaction, which lay the groundwork for further functional studies of miRNAs.

In light of chimera-defined miRNA:target pairs, we were able to globally assess the regulatory roles of miRNAs in the whole clock system consisting of input pathways, endogenous clock pacemaker and output pathways (Fig. 3). It is noteworthy that the disruption of *Clk* has a broad impact on the miRNA-mRNA interaction profile. The shifted pattern points to considerable changes of Ago1 RISC (RNA-induced silencing complex) assemblies in *Clk^jrk^*. One interpretation is that Ago1-miRNA loading is inefficient in *Clk^jrk^* flies because the *Clk* mutation could abolish some of the cyclic miRNA expression^53^. The miRNA-target interactions may also depend on the availability of miRNA targets, given the central role of *Clk* in circadian transcription regulation. The drastic changes of circadian mRNAs may alter the Ago loading or retention of the miRNA in RISC^61^. miRNAs regulating PDP1, VRI and TIM present distinct patterns, in which the miRNAs:*tim* interaction is *Clk*-dependent. Despite the different regulatory connection to *Clk* expression, the patterns may also result from the combinatorial regulation by multiple miRNAs with diverse cyclic expression patterns. The oscillating circadian mRNAs were regulated by multiple miRNAs to ensure correct timing of expression.

Our data clearly indicated that the 9 core circadian genes were regulated by 23 miRNAs in the brain of fruit flies (Fig. S3). Previous miRNA screens have identified a total of 47 miRNAs as relevant to circadian phenotype ^48–50^, 12 of those miRNAs were also detected in this study (Fig. S3D). The other miRNAs, which clearly have important roles in circadian rhythm, may indirectly regulate the expression of the circadian genes that have been confirmed to participate in circadian regulation^48–50^. In addition, we linked 10 miRNAs related to circadian rhythm and mapped 9 interaction sites on the *tim* transcript. These data suggest a complex regulation of miRNAs on rhythmic output.

The current study reports for the first time that *miR-375* is involved in circadian rhythm and sleep. At the larval stage of *Drosophila*, *miR-375* is expressed in the salivary glands and hindgut, but detailed expression characterization at adulthood is lacking^62^. In this study, we confirmed its circadian expression in the I-LNv neurons. Overexpressing *miR-375* with the *tim* driver leads to arrhythmic behavior with loss of morning anticipation and normal sleep pattern. Compared with an 81% arrhythmicity after overexpressing *miR-276a* in the previous study^47^, we observed a 100% arrhythmicity. In contrast to the peak expression of *miR-276a* at ZT22, the highest level of *miR-375* appeared at ZT12 with significant repression of TIM in the head at ZT18, as detected by immunoblotting (Fig.6). Multiple miRNAs may participate in the spatiotemporal control of *tim* expression, coordinating the normal rhythmic output.

Furthermore, we also found that overexpression of the *mir-305* in *tim* neurons (*iso:tim-gal4×UAS-luc-mir-305*) caused a unimodal activity and a phase shift in DD condition, while bimodal activity was maintained in an LD condition (Fig. S5). This phenotype was also observed in *miR-124* mutant^55, 56^. Our CLEAR-CLIP data suggest that *miR-124* targets *shaker (sh) and twenty-four (tyf)*. The chimera-defined miRNA-target interactions imply a broad impact of miRNAs on many aspects of the circadian system and provide a reliable guide to study the function of individual miRNAs. Extensive investigation is required to comprehensively understand the post-transcriptional regulation of circadian rhythms. This study opens a new avenue for a deep understanding of the post-transcriptional regulation of circadian rhythms.

## Acknowledgments

We thank Jeffery Price (University of Missouri at Kansas City) for revision of this manuscript. The work was supported by the Natural Science Foundation of China Grants 31572317 and 31730076 (to ZZ) and National Institutes of Health R01 AI122743 (to JZ).

## Author contributions

J.Z. and Z.Z. designed research; X.X., B.W. and X.F. performed research; X.X., X.F. analyzed data; and X.X., X.F., J.Z. and Z.Z. wrote the paper.

## Materials and Methods

### Fly strains

Fly strains were maintained on standard molasses-cornmeal-yeast food in a 12L:12D cycle at 25°C and 60% humidity. The fly lines were used in this study as follows: *w^1118^*, *Clk^Jrk^*, *USA-LUC-mir-305/TM3*, *UAS-LUC-mir-275.mir-305*, *UAS-LUC-mir-375*, *UAS-LUC-mir-9c*, *UAS-Mcd8::GFP*, *TI{GAL4}mir-375*[*KO*], which were purchased from the Bloomington Drosophila Stock Center. The *iso:tim-gal4* was obtained from Yirao’s lab*, UAS-tim2-5* from Jeffrey L. Price’s lab and was originally generated by Amita Sehgal’s lab.

### Ago1 CLEAR-CLIP library construction

The protocol was adapted from previous reports ^24, 39^. The collected tissue samples were first ground in liquid nitrogen before ultraviolet irradiation. The purified Ago1-RNA complexes from *Drosophila* lysates were subjected to stringent wash for removal of nonspecific binding. T4 RNA ligase 1 was added into the complex to promote the formation of miRNA-target chimeras. To track the size of the Ago1-RNA complex, RNA was labeled through ligation with a ^32^P-labeled 3’ DNA linker. A subsequent size selection on an SDS polyacrylamide gel was used to isolate the Ago1-miRNA-mRNA complexes (>130 kDa) and recover authentic target RNAs for library construction and sequencing.

### Behavior assay and analysis

Adult male flies (2-5 d old) were used to test locomotor activity rhythms. Flies were entrained under LD for 3d and released into constant darkness (DD) for at least 6 d at 25°C. Locomotor activity was recorded with *Drosophila* activity monitors (Trikinetics). Sleep was analyzed by pysolo and GraphPad software. FaasX software was used to analyze behavioral data. Locomotor behavior was analyzed in MATLAB. The details for the experimental protocol and data analysis were described by Chen et al^44^.

### Confocal microscopy

Adult male flies of 5-10 days were collected and their brains were dissected in phosphate buffered saline (PBS). The brains were subjected to 4% paraformaldehyde in PBS for 1 h and then washed three times with the wash buffer (0.05% Triton X-100 in PBS) for 10 min at room temperature. Samples were transferred to Blocking buffer (2% Triton-100, 10% normal goat serum in PBS) for an overnight incubation at 4 °C and incubated with primary antibodies (diluted in blocking buffer) overnight at 4°C. The following primary antibodies were diluted in the blocking buffer: rat-anti-TIM (1:1000, from Jeffrey L. Price), mouse-anti-PDF (1:400, from DSHB). After washing samples three times for 15 min at room temperature, the samples were incubated with secondary antibody (1:200) at 4°C overnight. The samples were imaged on a Leica SP8 confocal microscope. ImageJ software was used for TIM and PDF quantification.

### RNA extraction and qRT-PCR analysis

Fly heads were collected at the indicated time points and stored at −80°C before RNA extraction. Total RNA was extracted from the heads by using Trizol reagent (TIANGEN) following the supplier’s instruction. For reverse transcription and real-time PCR of *timeless*, we used PrimeScript RT reagent kit with gDNA Eraser (TAKARA) and superreal premix plus (SYBR Green) (TIANGEN). The sequences of primers are shown in Table S4. For quantitative analysis of *timeless*, *miR-375* and 2s rRNA, we used a miRcute miRNA first-strand cDNA synthesis kit and a miRcute miRNA qRT-PCR detection kit (SYBR Green) (TIANGEN). The miRNA-specific forward primers used for qRT-PCR are also shown in Table S2.

### Western blot

Flies’ heads were collected at the indicated time points and homogenized in a 1.5 mL microtube with RIPA lysis buffer (strong) supplemented with proteinase inhibitors. For immunoblot analysis, proteins were transferred to PVDF membrane. After blocking, the membrane was incubated with actin antibody (1:10000) and TIM antibody (1:4000) for 2 h at room temperature. Image J was used to calculate band intensity.

### Identification of miRNA-mRNA chimeras

Ago1 CLEAR-CLIP sequencing reads were preprocessed to remove low-quality reads and 3’ adapter sequences using Flexbar^63^. Shorter reads (less than 16 nt) were discarded. Reads with identical sequences were collapsed. Three random nucleotides were used in the 5’ adapter to avoid PCR over-amplification. These random barcodes were trimmed prior to mapping.

The clean reads containing miRNA sequences were first defined by mapping mature miRNA sequences against sample libraries using BLAST (v2.3.0) with e-value equals to 0.4^64^. Only the best alignment was reported. The remaining sequences excluding miRNAs part were extracted for mapping to the transcripts with BLAST. The reads mapped to rRNA, tRNA and miRNA genes were removed.

### Peak calling of CLIP cluster

The peak calling of CLIP cluster was done as previously described using the pooled reads from three biological samples at each time point^65^. Briefly, overlapped reads were collapsed to form cluster regions. Cubic spline interpolation (Scipy, http://www.scipy.org/) was performed to determine the location and read number of peaks within a cluster^66^. Significant peaks were identified by determining reads number cutoffs with the p-value less than 0.01 using Poisson distribution.

### Motif analysis

The analysis of overrepresented motifs of Ago1 CLEAR-CLIP by MEME was performed as previously described^59^ with the settings: -dna -mod zoops -max w7 -n motifs 1. The motifs identified by meme for targets of each miRNA were then aligned to the reverse-complemented miRNA sequence using FIMO^68^, with the setting-output-p thresh 0.01. High-confidence motifs were generated with FIMO q-value (FDR) < 0.05, and meme Bonferroni-corrected p-value < 0.05.

For *de novo* motif analysis using Homer, miRNAs with more than 30 unique chimeras-defined sites were kept. The background sequences were randomly selected from other miRNA chimeras. Information score (c) was calculated as previously described^59^ to retain motifs with c≥1. Heatmap with the percentage of the motif presence was created using R heatmap package.

### RNA duplex structure prediction and binding free energy

The predictions of duplex structure for miRNAs and their target sequences were performed by RNA hybrid. Mutations in chimeras sequences for both miRNA and mRNA were omitted by comparing with the original annotation. The target sequences were extended to 100 nt according to the miRNA seed sites distribution in *D. melanogaster*. The setting of RNA hybrid was modified as previously described to favor the seed pairing^59^. The minimum free energy of each interaction was calculated by RNA hybrid with default settings^67^. The shuffled interactions served as controls for the comparison of free energy distribution. The median free energy for each profile was calculated in R for comparison.

The duplex heatmap was performed with transformed pairing data (base pairing including G-U match as 1, unpaired site as 0). Cluster numbers (k) 3-10 were tested, with k=5 providing the most meaningful set of distinct categories.

### Analysis of chimera targets in miRNA perturbation experiments

The gene expression data after miRNA perturbation was downloaded^45, 46^. The log2 fold-change was used to perform cumulative distribution function (CDF) analysis comparing transcripts bearing chimeras-derived miRNA target sites versus those without target sites. The CDFs were also plotted for transcripts in which the chimera-derived miRNA target sites overlap with Ago1 CLIP peaks.

### Dynamic analysis of miRNA-mRNA interactions

To quantitatively analyze the temporal profile of miRNA-mRNA interactions, the chimera-supported Ago1 binding peaks were normalized to the total unique reads in each condition^69^. In order to visualize the pattern of difference between each time point, the data were loaded into Gene Expression Dynamics Inspector (GEDIv2.1) with default parameter (26×25 grids) to perform the dynamic analysis^70^. GEDI employs the SOM algorithm to assign the interactions with similar trend into close tiles. To select the interaction cluster presenting distinct pattern at a certain time point, we first located the tile with maximum value at local spots displaying unique interactions and expanded the spots following gene Density map in GEDI to the edges. Unique interactions for each category were extracted for phenotype enrichment analysis based on FlyBase annotation.

### GO and KEGG enrichment analysis

The selected gene list was taken as an input for the functional enrichment analysis with DAVID^71^. The top-ranked GO terms belonging to biological functions were chosen.

### Statistics analysis

Statistics analysis for all indicated data in this study were performed with T-test and p values considered significant at p < 0.05 and extremely significant at p < 0.001.

---

**Fig. S1.**
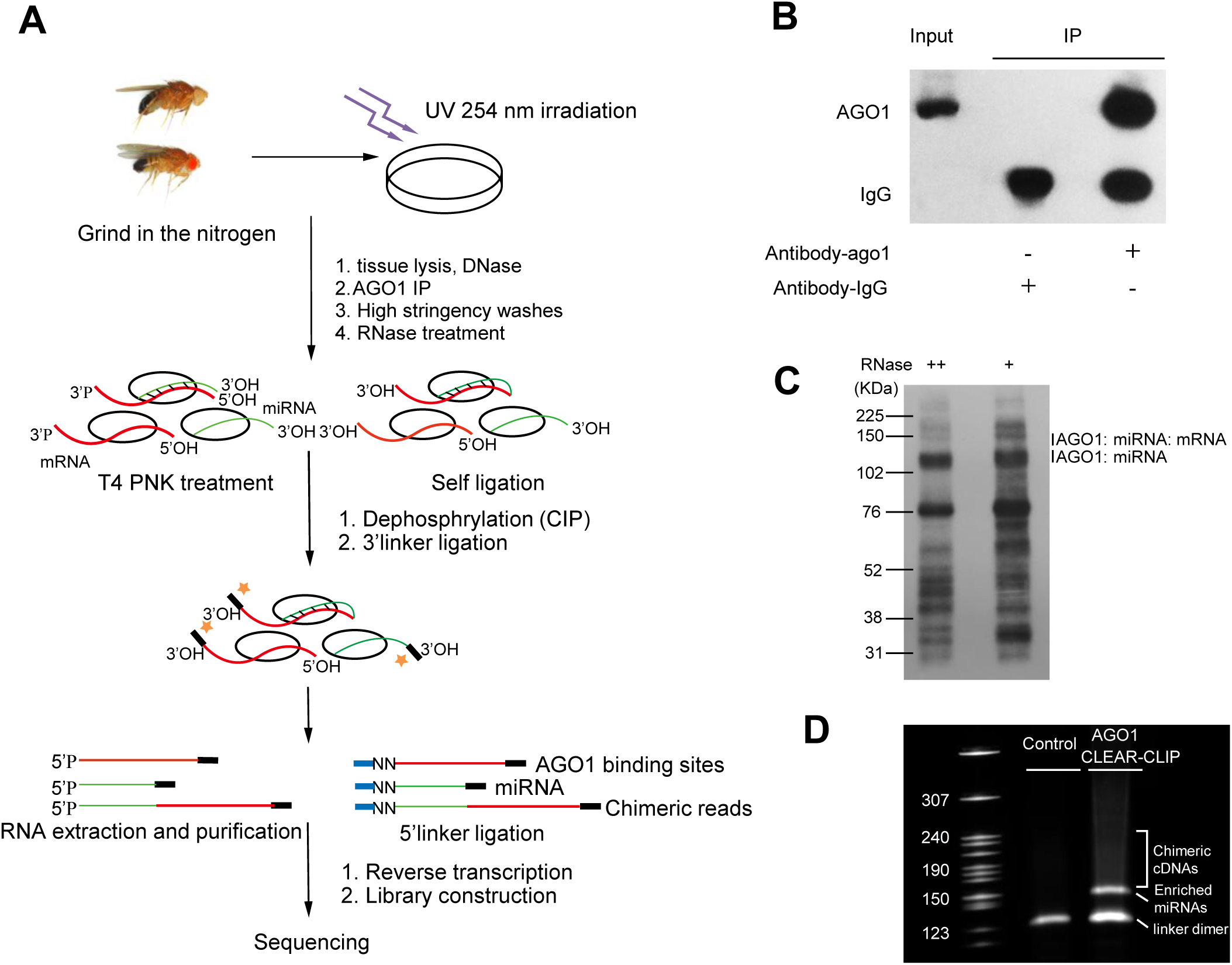
Experimental Procedure of Ago1 CLEAR-CLIP. (A) In Ago1 CLEAR-CLIP, the tissue lysates were prepared from UV cross-linked flies. Endogenous Ago1 was immunopurified and stringently washed to remove non-specific interactions. The RNA ends in the Ago1-complexes were treated with T4 Polynucleotide Kinase and ligated together. RNAs associated with Ago1 were then further purified by size selection using the SDS-PAGE gel and recovered for library construction and sequencing (material and method). (B) Western blot detect Ago1 in input sample, IP with antibody-Ago1 and antibody-IgG. (C) Autoradiogram of the Ago1 associated RNA after digestion with different RNase. (D) PCR products amplified using the recovered RNAs after linker ligation.

**Fig. S2.**
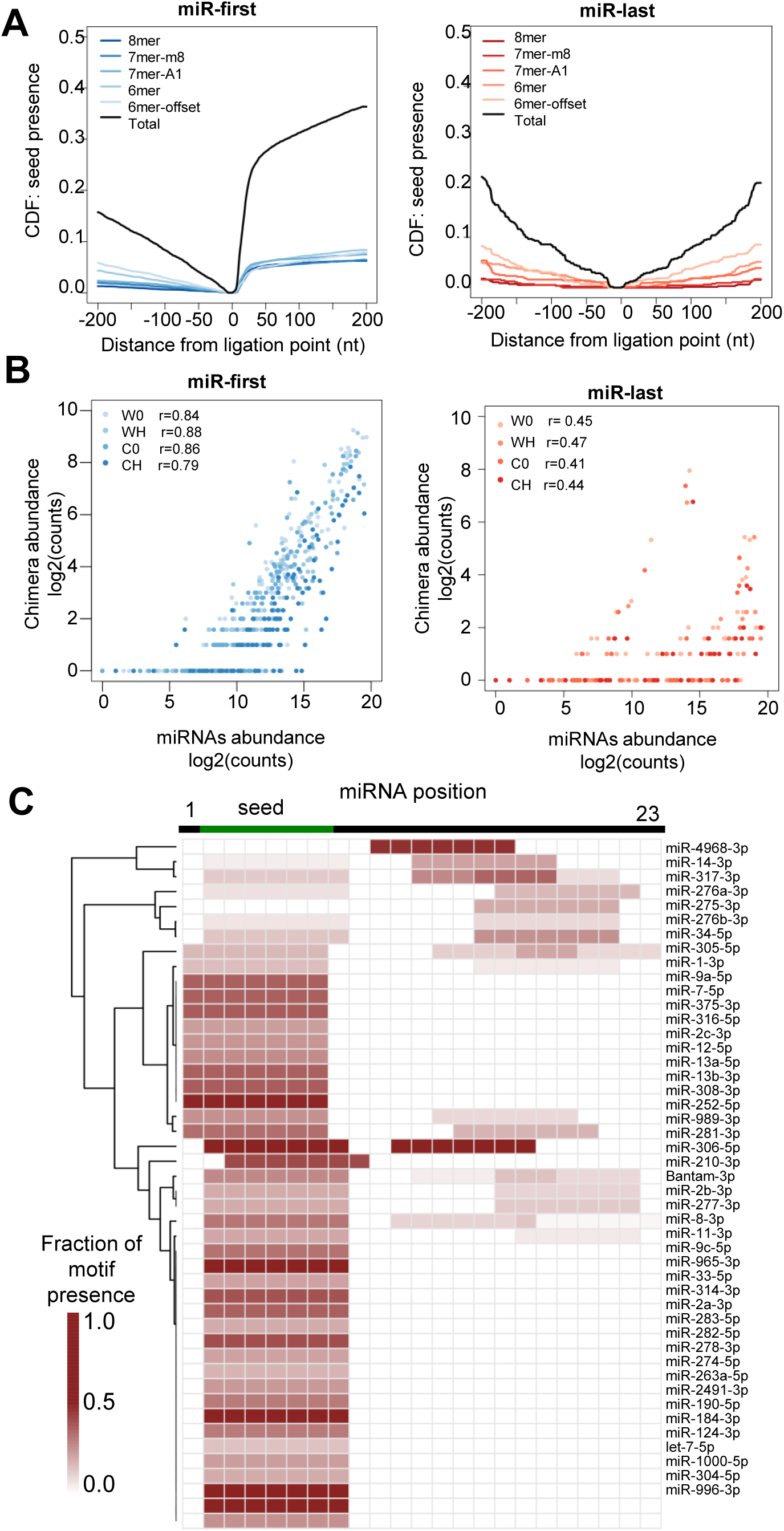
miRNAs detected in the Ago-RNA complexes and the miRNA-mRNA chimeras. (A) The cumulative distribution function of canonical miRNA seed matches in target regions relative to ligation site for miR-first and miR-last chimeras. (B) Correlation plots of miRNA abundance from the whole CLEAR-CLIP libraries and their chimeras. Pearson’s correlation coefficients are shown for each condition. (C) The heatmap shows the distribution of significant enriched 7mer motifs through de novo analyzing all chimeras detected for individual miRNA. The colour intensity is proportional to abundance of 7mer motif in target sequences. The hierarchical clustering was performed in R package gplots2.

**Fig. S3.**
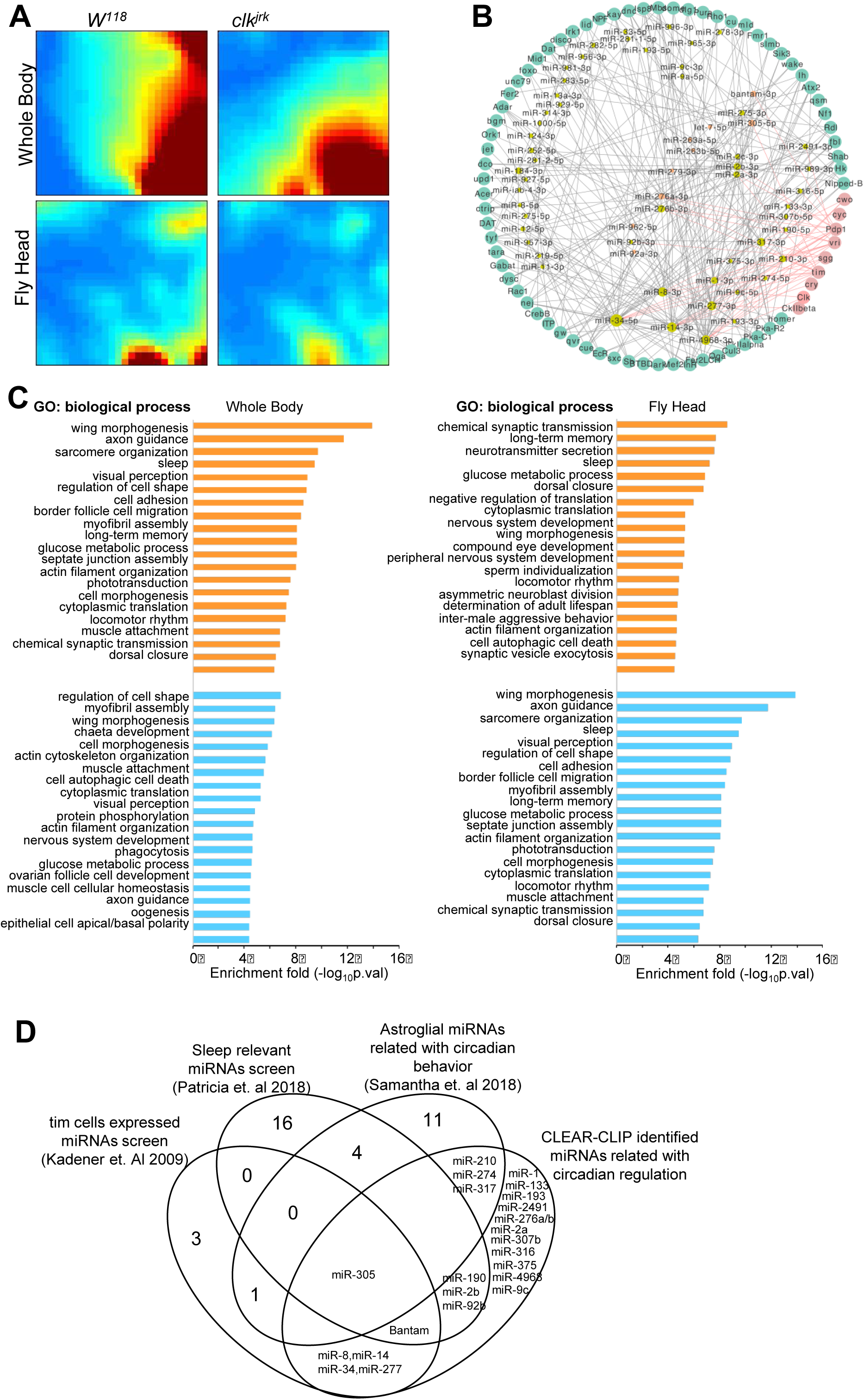
The miRNA-mRNA interactions involved in Drosophila circadian system. (A) Globally distinct miRNA-mRNA interactome of wild type W1118 and Clkjrk mutant in whole body or head displayed as self-organized maps with the GEDIv2.1. Color bar indicates the normalized peak height. (B) The miRNAs:circadian-genes interacting networks defined chimeras from *Drosphila* CLEAR-CLIP. The circadian genes were extracted from Flybase with phenotype emphasized in circadian defective behavior. These genes are aligned in circle, color in light red represents core genes in circadian system, color in light green stands for other circadian related genes. miRNAs in red have published results supporting circadian related functions or expression. (C) GO enrichment analysis of chimeras-defined miRNA targets. The top twenty biological process terms were plotted. (D) The Venn diagram of circadian-relevant miRNAs screen.

**Fig. S4.**
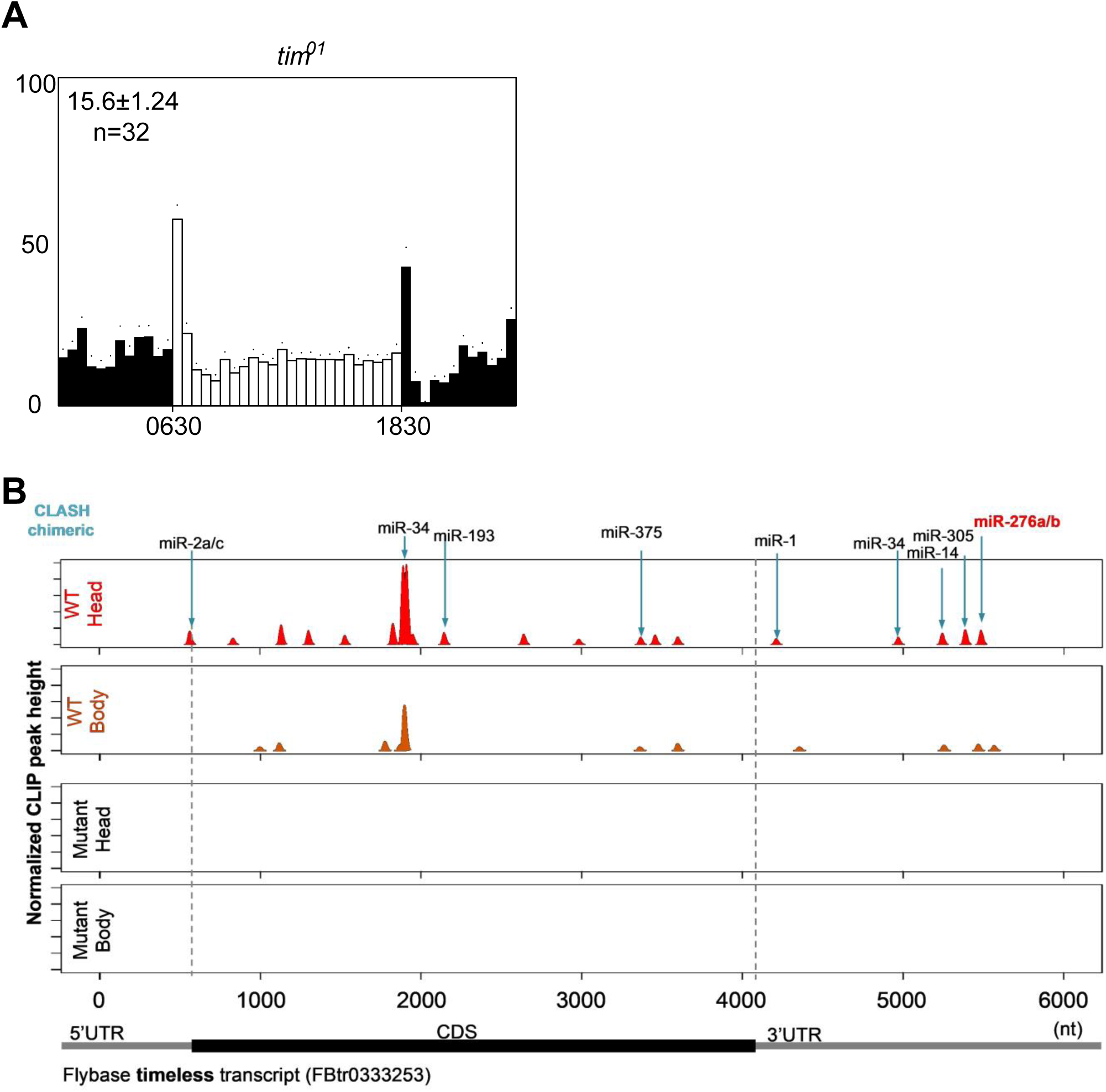
The miRNAs involved regulation of timin *Drosophila*. (A) Locomotor profiles of tim01 in LD condition. (B) The distributionofAgo1 peaks along the timeless transcript. miR-276a has been confirmed to target timelss.

**Fig. S5.**
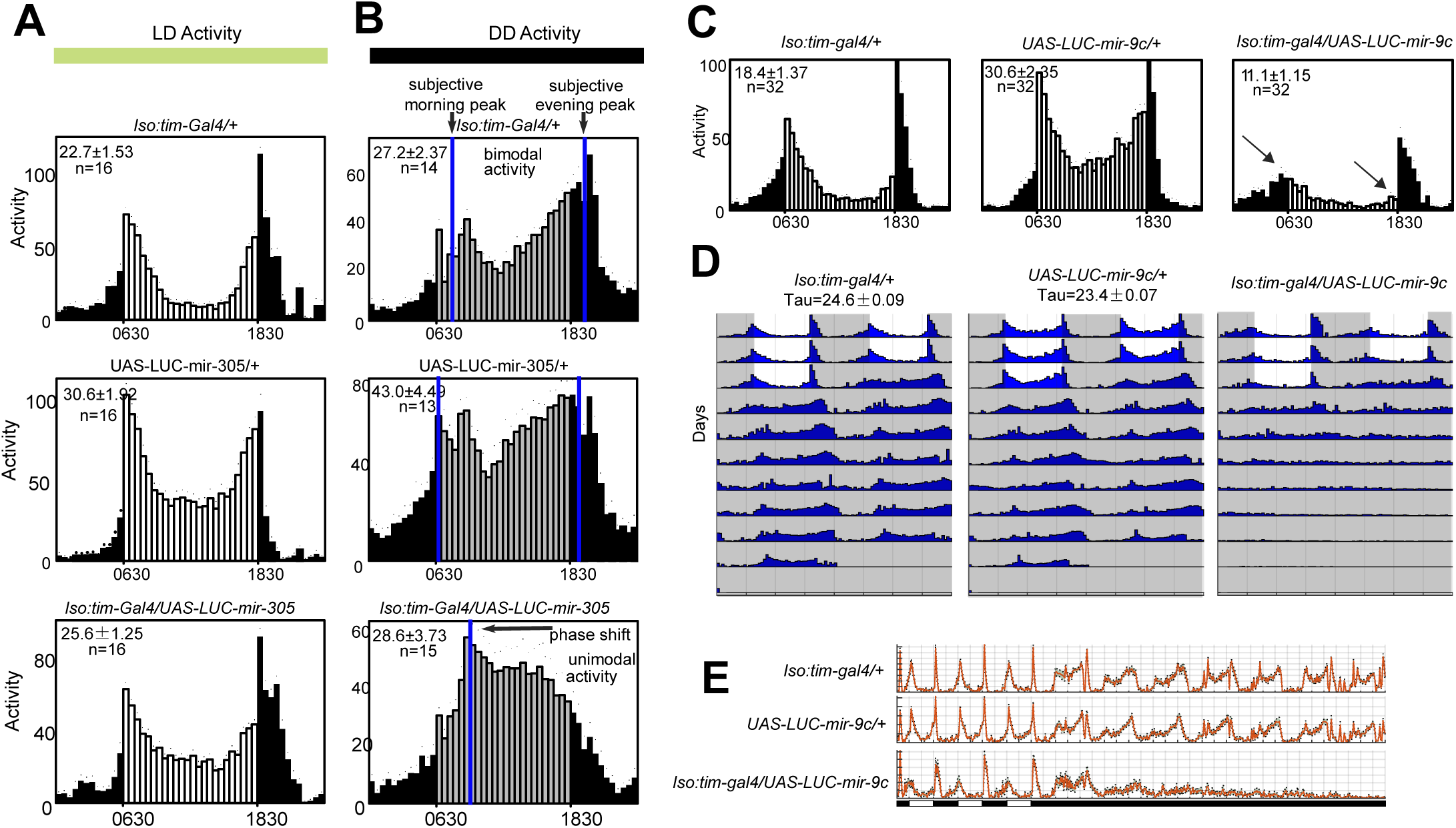
Overexpress mir-305/mir-9c in tim-neuron affects phase of circadian locomotor behavior. (A) Average activity profiles across three light/dark cycles, mir-305 OE flies (iso:tim-gal4/UAS-LUC-mir-305) exhibit normal bimodal profiles just like controls (iso:tim-gal4/+; UAS-LUC-mir-305/+). (B) Average activity profiles during the first two DD cycles follow transfer to DD. Controls exhibit a bimodal DD activity profile with peaks around subjective morning and evening. Overexpression of mir-305 induces a unimodal profile in which evening peak activity is advanced. (C) Comparison of the circadian locomotor activity in light/dark cycles of controls (iso:tim-gal4/+;UAS-LUC-mir-9c/+) and mir-9cOE (iso-tim-gal4/UAS-LUC c) flies. Histograms represent the distribution of activity through 24h, averaged for flies over three LD days, morning and evening anticipation are lost in mir-9c OE flies (dark raw). (D) Locomotor behavior under LD cycle and constant darkness. Representative double plotted actograms of *iso:tim-gal4/+, UAS-LUC-mir-9c/+, iso:tim-gal4/UAS-LUC-mir-9c* flies. White indicates the light phase, gray indicates the dark phase. (E) Activity analysis of flies with overexpression of *mir-9c* driven by iso-tim-gal4 in 3LD and 7DD.

**Table S1A.**
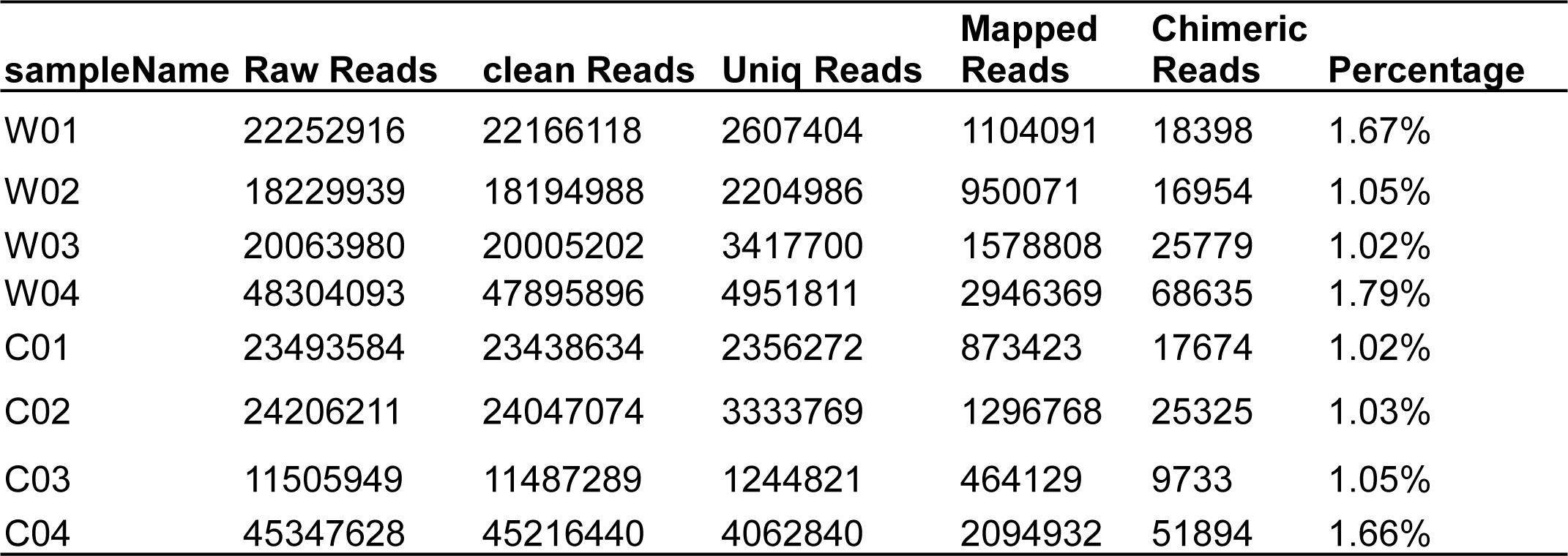
Read statistics for 8 drosophila CLEAR-CLIP samples.

**Table S1B.**
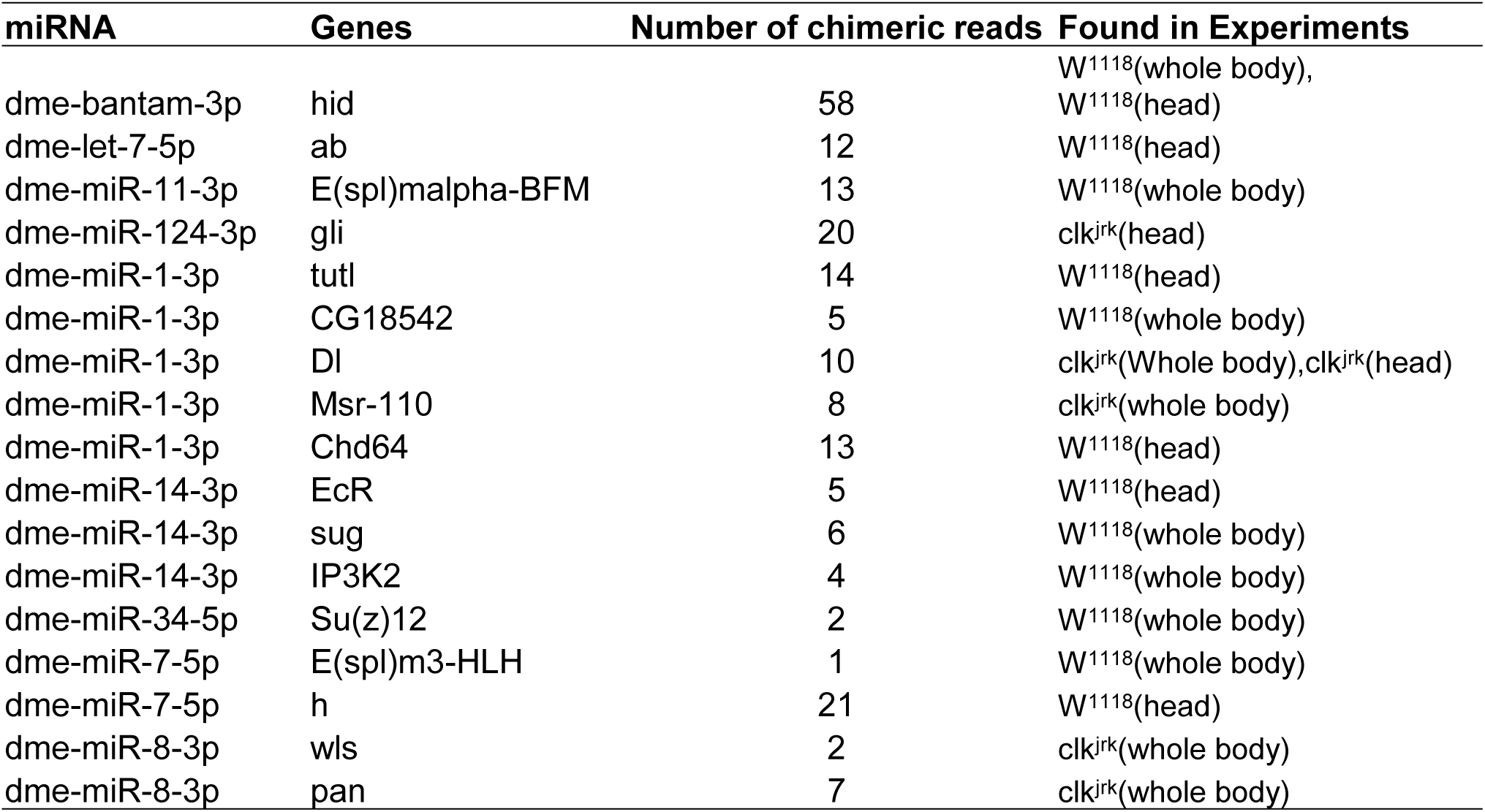
Experimentally Validated miRNA Targets Found in chimeras supported interactions.

**Table S1C.**
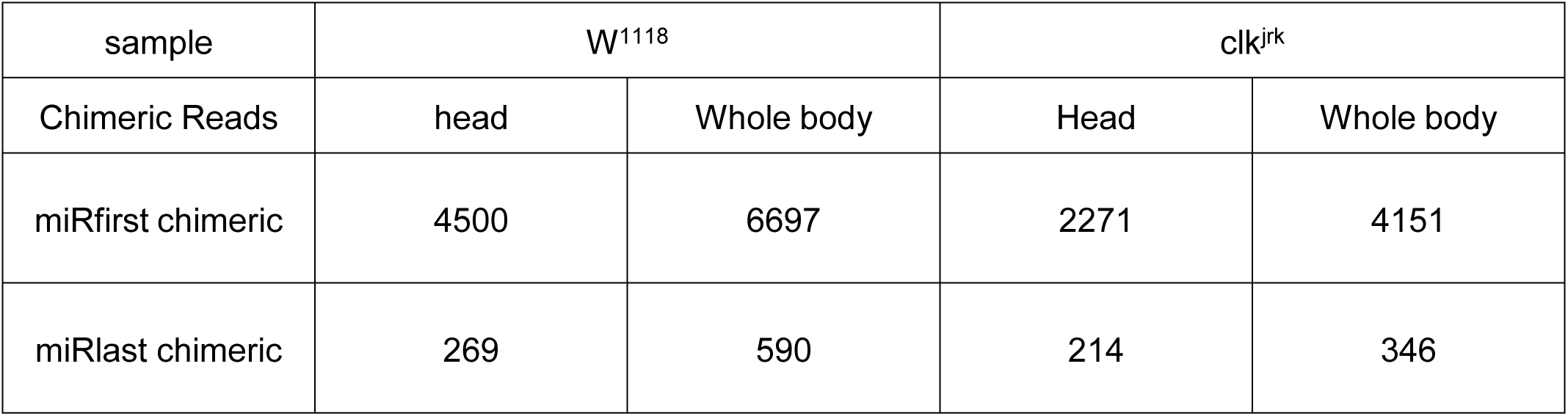
miRNA-first and miRNA-last distribution found in CLEAR-CLIP.

**Table S2.**
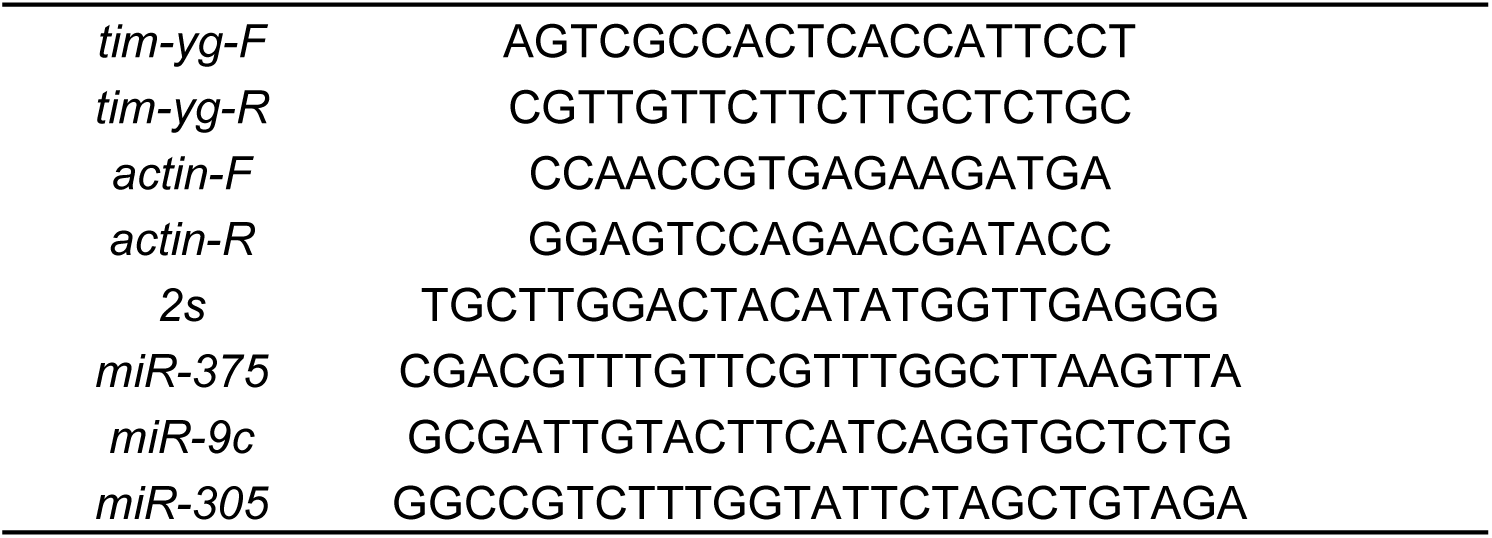
The sequences of primer for real time PCR.

